# Improvement of baculovirus as protein expression vector and as biopesticide by CRISPR/Cas9 editing

**DOI:** 10.1101/662890

**Authors:** Verónica Pazmiño-Ibarra, Adriá Mengual, Alexandra Targovnik, Salvador Herrero

## Abstract

The CRISPR (Clustered Regularly Interspaced Short Palindromic repeats) system associated Cas9 endonuclease is a molecular tool that enables specific sequence edition with high efficiency. The edition using CRISPR/Cas9 system has been successfully reported in small and large viral genomes. In this study, we have explored the use of CRISPR/Cas9 system for the edition of the baculovirus genome. We have shown that the delivering of Cas9-sgRNA ribonucleoprotein (RNP) complex with or without DNA repair template into Sf21 insect cells through lipofection might be efficient to produce knocks-out as well as knocks-in into the baculovirus. To evaluate potential application of our CRISPR/Cas9 method to improve baculovirus as protein expression vector and as biopesticide, we attempted to knock-out several genes from a recombinant AcMNPV form used in the baculovirus expression system as well as in a natural occurring viral isolate from the same virus. We have additionally confirmed the adaptation of this methodology for the generation of viral knocks-in specific regions of the viral genome. Analysis of the generated mutants revealed that the edition efficiency and the type of changes was variable but relatively high. Depending on the targeted gene, the rate of edition ranged from 10% to 40%. This study established the first report revealing the potential of CRISPR/Cas9 for the edition of baculovirus contributing to the engineering of baculovirus as protein expression vector as well as a biological control agent.

## INTRODUCTION

Baculoviruses are rod-shaped DNA viruses with double stranded circular genome that infect invertebrates, particularly insects of the order Lepidoptera (1). They are widely used as biopesticide as well as versatile and powerful vector for recombinant protein expression (2). In nature, the baculovirus mainly infects Lepidoptera larvae when they feed on plant contaminated with the virus (3). During their viral biphasic life cycle, baculoviruses produce two distinct virion phenotypes: occlusion derived viruses (ODV) and budded viruses (BV). While ODV are involved in horizontal virus transmission from insect to insect through structures named occlusion bodies (OB) which have the virus embedded within, BV are involved in spread of the infection from cell to cell (4).

Autographa *californica multiple nucleopolyhedrovirus* (AcMNPV), the prototypic baculovirus most common used for biotechnological purpose, has a circular genome of about 134 Kb that contains 156 predicted open reading frames (ORF) (5). The baculovirus genomes, and specially AcMNPV genome, have been genetically modified in order to enhance their pesticide potency and increase the quality and quantity of the recombinant protein expressed in the system (6, 7). These modifications include either the deletion of non-essential genes for virus survival or infectivity and the insertion of foreign genes. For instance, in term of increase the insecticidal activity of baculovirus, the cry1Ab gene from *Bacillus thuringensis* and neurotoxins from scorpion venom have been incorporated into the baculoviru*s* genome (8, 9). In addition, the *egt* gene (*Ac15*), involved in the inhibition of the molting, has been deleted from the virus significantly improving the speed of killing of the virus (10). Moreover, different attempt has been conducted to improve the heterologous protein expression in the baculovirus expression system (BEVS). It has been reported that simultaneous deletion of non-essential genes for the *in vitro* replication of AcMNPV can enhance the expression of recombinant proteins (11, 12). Another strategy implemented has been the addition of heterologous genes into de viral genome with valuable properties for protein production (7).

The baculovirus expression vector system (BEVS) has become a powerful platform to produce large quantity of recombinant proteins both in insect cells (Sf21, Sf9 and High Five™ cells) as in Lepidoptera larvae. The recombinant virus was originally obtained by replacing the polyhedrin gene from AcMNPV by co-transfection in insect cells with a viral genome and a donor plasmid (13). Different improvements have been developed to optimize the method for the generation of recombinant baculovirus. A remarkable success was the construction of the first bacmid that contained the whole AcMNPV genome and could can be propagated in *Escherichia coli* cells (14). The system, commercialized as Bac-to-Bac® system, is based on the introduction of a heterologous gene from a shuttle vector to the bacmid by transposition, producing a 100% recombinant progeny.

In the last years, multiple approaches for viral genome engineering have been develop. Among the diverse genome editing technologies available, the clustered regularly interspaced short palindromic repeats-Cas9 system (CRISPR/Cas9) appears as the most successful tool for editing large viral genome (15). The system consists of a Cas9 endonuclease from *Streptococcus pyogenes* that cleaves double-stranded DNA (dsDNA) and an RNA complex (sgRNA) that direct sequence-specific dsDNA cleavage by Cas9. The double-strand break (DSB) are recognized and repaired by the cell endogenous mechanisms through non-homologous end joining (NHEJ) that produce insertions or deletions (indels), or homology-directed repair (HDR) in presence of a donor template DNA with homology to the sequence flanking the DSB (16–18). Recent developments in this technology have made possible to generated precise modifications into a wide variety of viral genomes (19). However, the CRISPR/Cas9 system has not been applied for the edition of baculoviral genomes.

In this study, we developed for first time a CRISPR/Cas9-assisted method to edit AcMNPV genome in multiple ways and with different purposes. We have shown that the delivering of Cas9-sgRNA ribonucleoprotein (RNP) complex through lipofection in insect cells might be efficient to generate gene knock-out and knock-in. To evaluate potential application of our CRISPR/Cas9 method, we attempted to knock-out several genes from a recombinant baculovirus used for the BEVS to study their effect on recombinant protein production as well as in a natural occurring viral isolate to assess the potential of the methodology to improve baculovirus as a pesticide. In addition, we have also show that our methodology can also be used for the efficient introduction of foreign DNA on the genome of baculovirus.

## RESULTS

### AcMNPV gene knock-out

By direct transfection of the Cas9/RNP complex together to the viral genome of AcMNPV we were able to introduce different mutations in the targeted genes. The procedure was reproduced in several non-essential genes such as the *ODV-E26, F-Protein, p74* and *Ac18* with potential application to increase the recombinant protein production using the BEVS. In addition, we also attempted to edit *egt* gene to enhance the insecticidal properties of wild type (wt) AcMNPV. The Cas9/sgRNA complexes were assembled *in vitro* for each target gene described in Table 1. Individual viral clones were isolated from each edition by plaque assay and presence of indels in each of the selected clones was assessed by PCR amplification of the targeted region and Sanger sequencing.

**Table 1.**
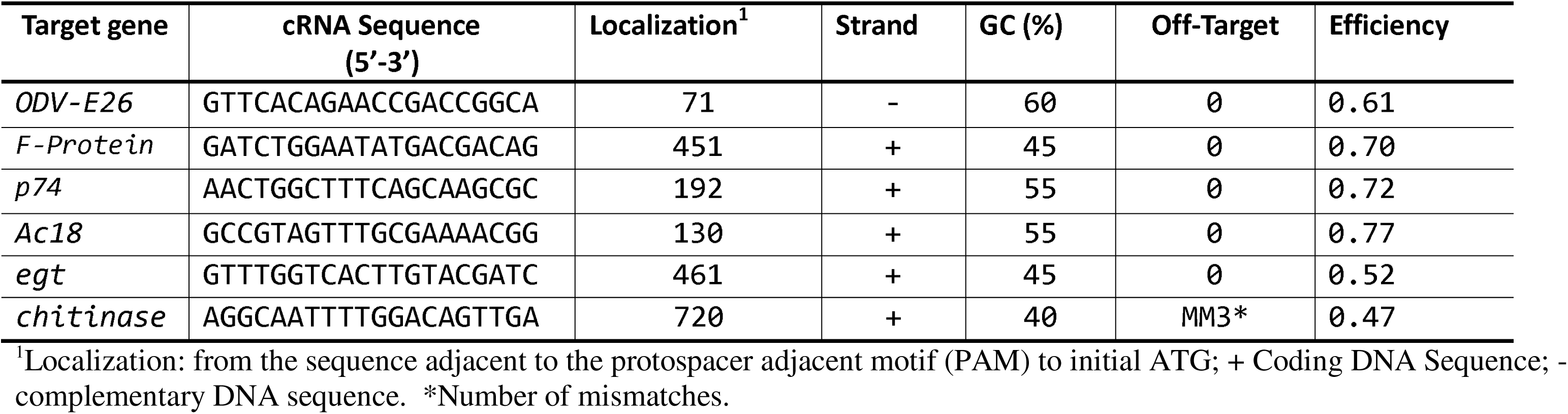
Summary of sgRNA characteristics

Sequencing analysis revealed the successful edition of several clones of each of the targeted genes (Table 2). The edition efficiency and the type of changes was variable. Depending on the targeted gene, the rate of edition ranged from 10% to 40% of the analyzed clones. All the edited viruses had a deletion on the targeted gene, except for the *F-Protein*#1 that showed an insertion of 125 nucleotides and *ODV-E26*#1 that contains a deletion of 4 nucleotides and an insertion of 16 nucleotides (Table 2). Except for one clone (*ODV-E26*#1), all the edited genes had a deletion located at the expected region (adjacent to the PAM site). In agreement with the strand targeting by the sgRNA (Table 2), all the deletion at the ODV-E26 gene were located down-stream from the PAM sequence, while deletion for the other genes were located up-stream from the PAM sequence. The effect of edition on the predicted protein was diverse. The different editions introduced frameshift mutations producing early stop codons (*ODV-E26#2-4, F-Protein*#1, *p74*#1, *Ac18*#1, and *egt*#2-4) or produce changes and deletion of few amino acids in the predicted protein (*ODV-E26#1, Ac18*#2, and *egt*#1) (Table 2).

**Table 2.**
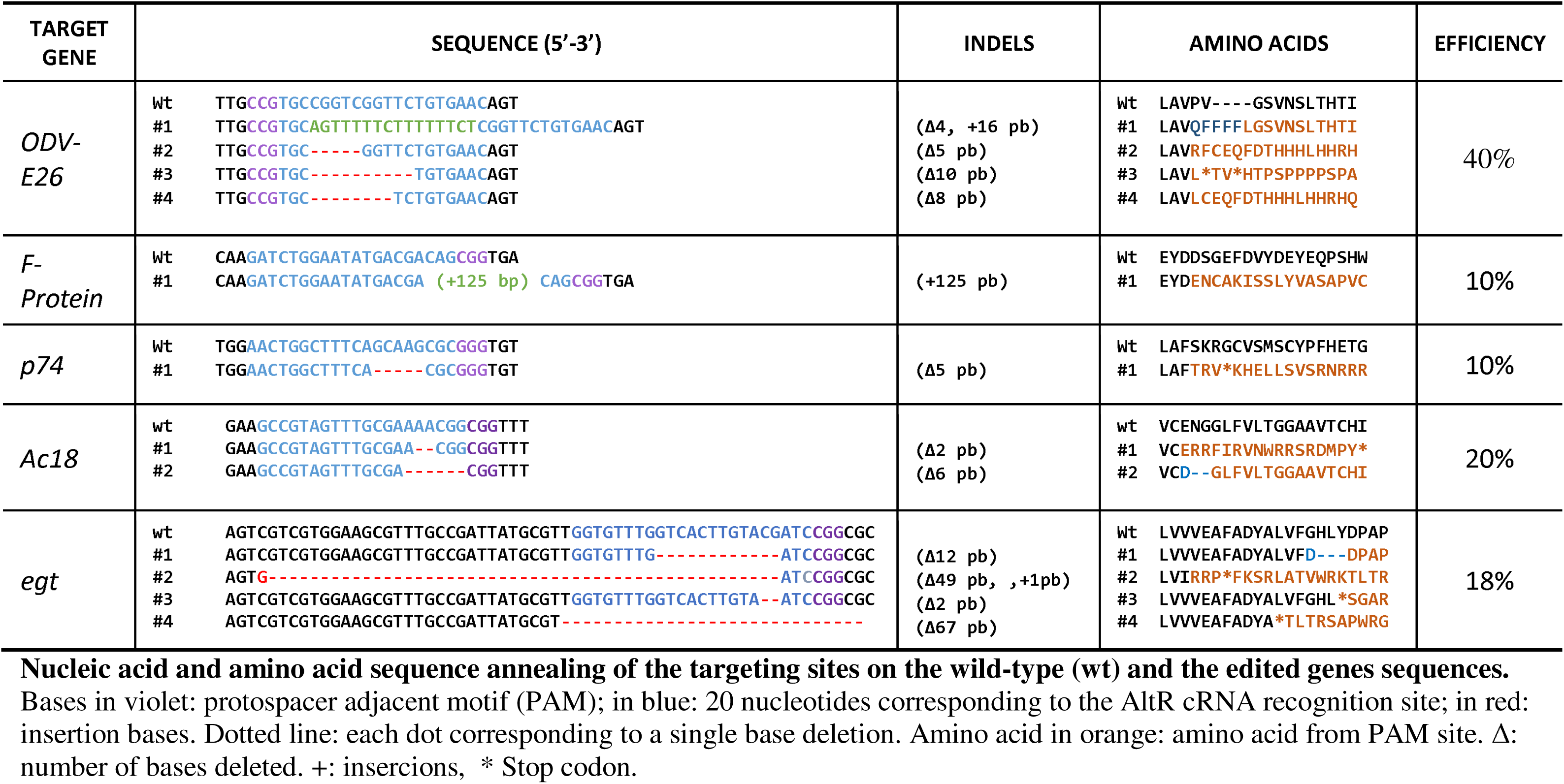
Summary of sequencing analysis of indels in target-site

### Effect of mutations on viral replication and recombinant protein production

To compare the performance of the individual viruses in cell culture we focused on the mutants with a stop codon or a frameshift mutation derived from the edition of the recombinant virus (derived from Bac-to-Bac® system) and expressing the GFP protein under the *pSeL* promoter. No significant differences in the production of budded viruses was found among the different viruses, suggesting that the produced knocks-out were not affecting the viral infectivity and replication in cultured cells (Fig. 1). Next, we tested the recombinant protein production in Sf21 cell of the different viruses. For this purpose, Sf21 cells were infected with the AcMNPV mutants at MOI 0.5 (Fig. 2A) and 5 (Fig. 2B). Three days post-infection, the cells were harvested and the GFP activity was measured. No statically significant differences between the control and the mutants were found (p-value > 0.05) suggesting that knock-out of those genes does not have major effect on the recombinant protein production in cell culture.

**Figure 1.**
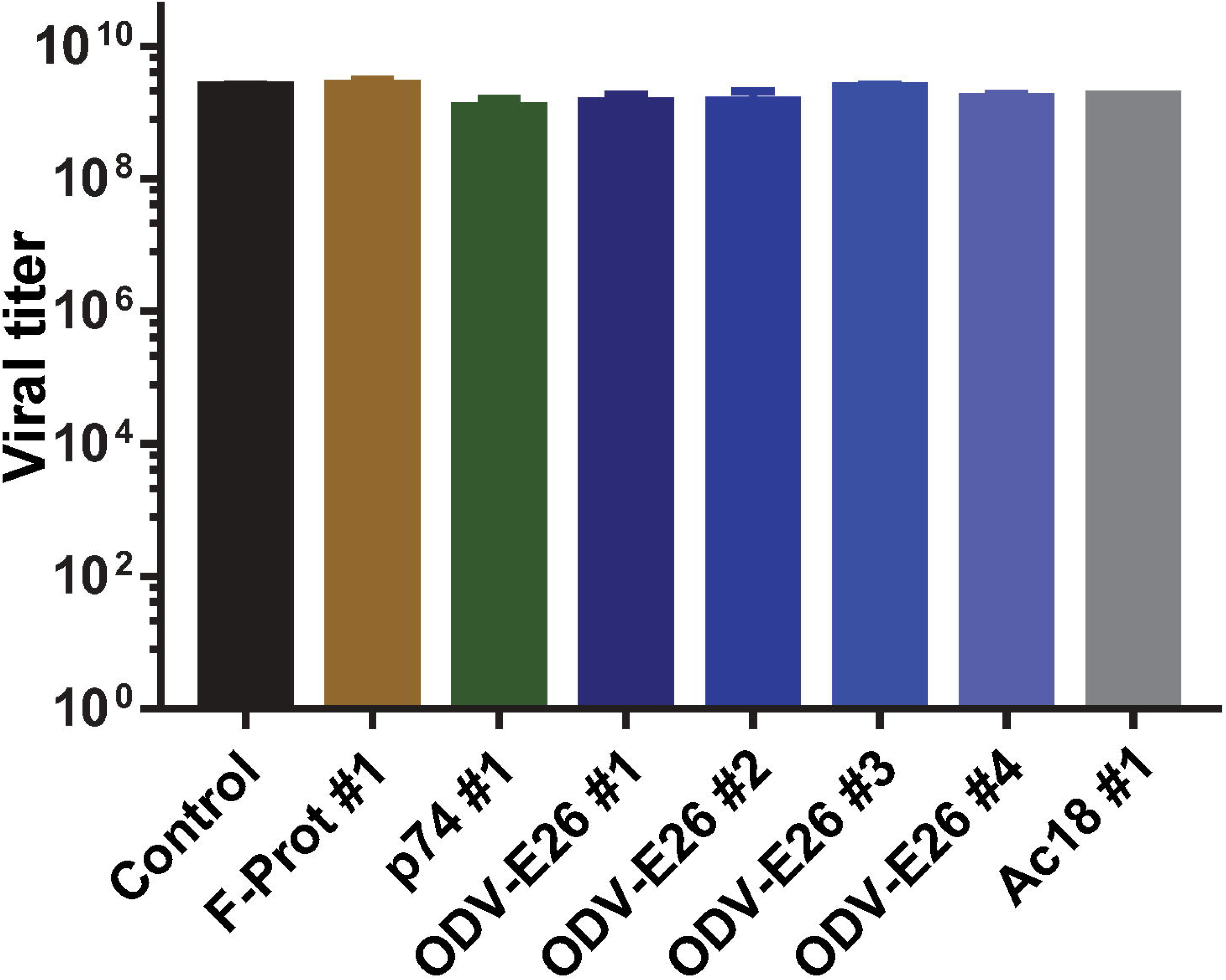
Effect of mutations on viral replication. Final viral titer of the different AcMNPV mutants after 72 hours post infection in Sf21 cells infected at MOI 5. A non-edited virus was included as a control.

**Figure 2.**
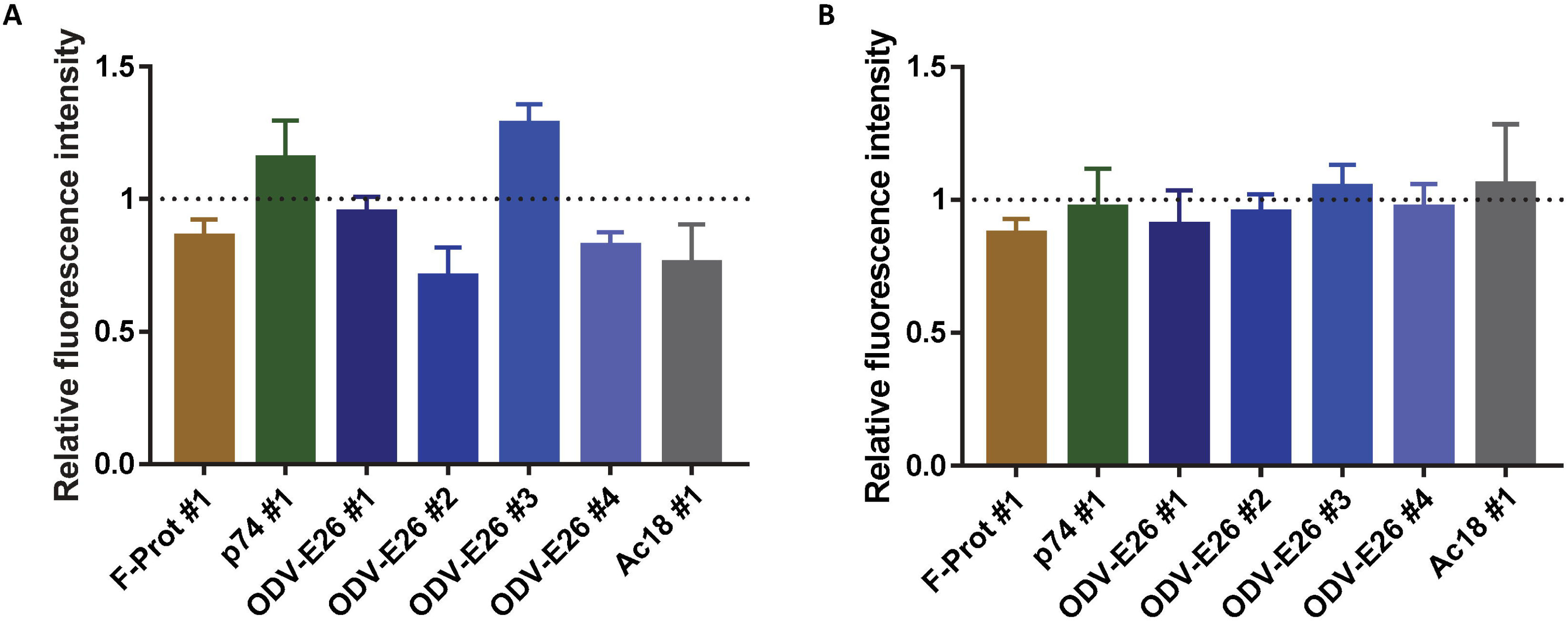
Effect of mutations on recombinant protein production in cell culture. GFP expression level in Sf21 cells infected with the different edited AcMNPV at MOI 0.5 (A) and 5 (B). The GFP expression was measured as relative fluorescence intensity at 72 hours post infection. The results are expressed as the relative GFP fluorescence intensity, taken as 1 of the value corresponding to the maximum intensity obtained with the control virus. The values are the means of at least three independent assays. The error bars represent the standard error of the mean. A non-edited virus was included as a control.

Insect larvae can be used as an inexpensive alternative to fermentative technologies for protein production (7). We tested the effect of the previous mutations in protein production in *S. exigua* larvae. Last instar larvae were infected by intrahemocelical injection with the AcMNPV mutants. 72 hours post-infection, GFP expression was estimated by measuring the fluorescence intensity from the cellular extracts of the larvae (Fig. 3). No differences in GFP production were observed for the F-protein, p74, and Ac18 knocks-out. Although, all the ODV-E26 mutants showed protein expression values above the control viruses, only the values for the ODV-E26 #1 and ODV-E26 #3 had statistically significant differences, with GFP production values of at least five times higher than larvae infected with the control virus. Simultaneous quantification of the different ODV-E26 clones together reported a significant increase of about 5-fold in protein production (Fig.3). These results revealed that simple knock-out of *ODV-E26* using the CRISPR/Cas9 can be used to increase the recombinant protein yield in insect larvae.

**Figure 3.**
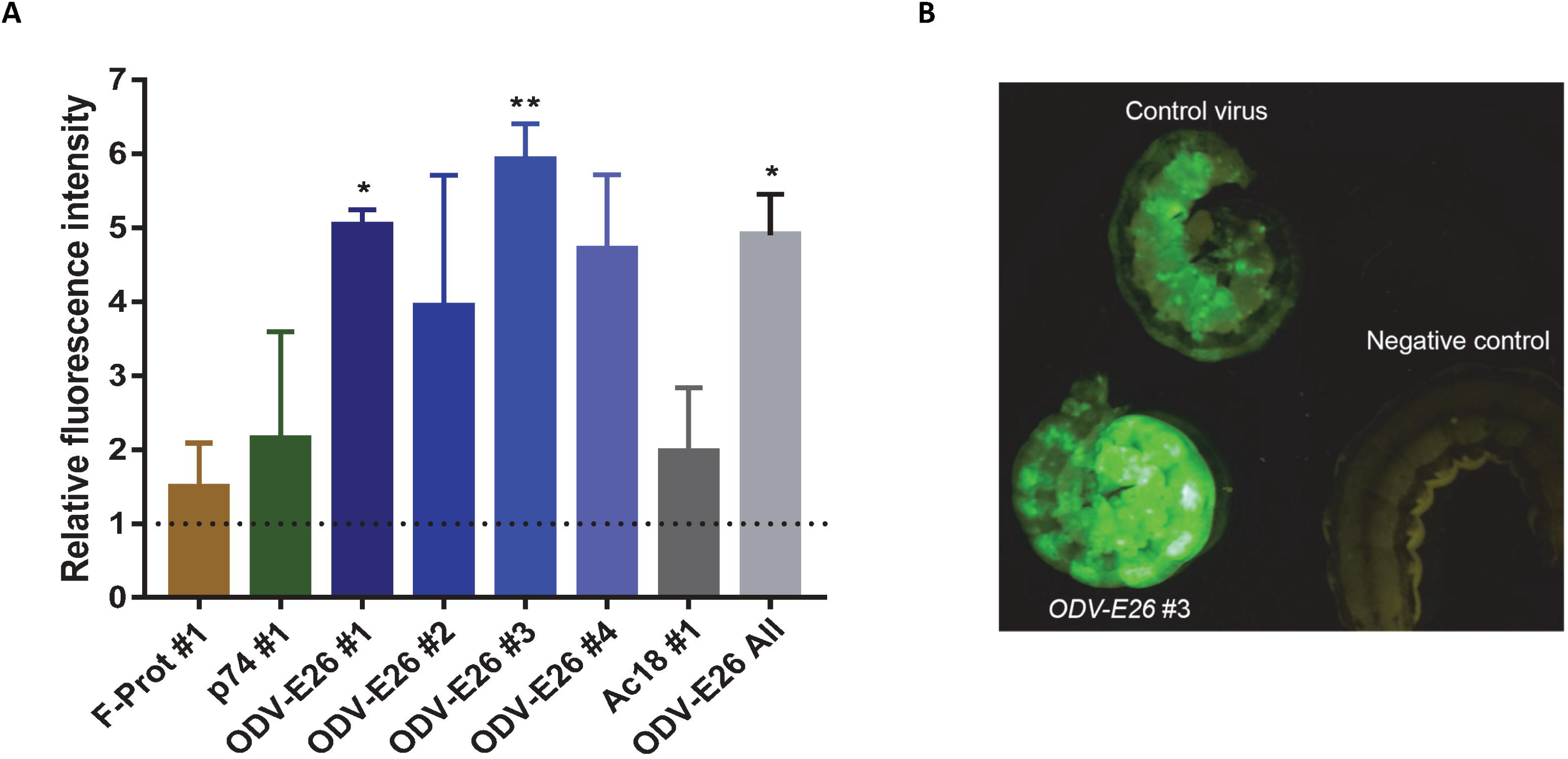
Effect of mutations on recombinant protein production in larvae. Analysis of GFP expression level in *S. exigua* larvae infected with the different edited AcMNPV (A). The GFP expression was measured as relative fluorescence intensity at 72 hours post infection. The results are expressed as the relative GFP fluorescence intensity, taken as 1 of the value corresponding to the maximum intensity obtained with the control virus. The values are the means of at least three independent assays. The error bars represent the standard error of the mean. A non-edited virus was included as a control. The average value obtained with the four different ODV-E26 mutants is also reported (light grey bar). Representative image of *S. exigua* larvae infected with the recombinant virus (72 hours post infection) (B).

### Insecticidal activity of AcMNPV mutant

We have additionally showed the generation of CRISPR/Cas9 knocks-out on the wild type AcMNPV. Next, we tested if such viruses could become a recombinant free alternative to improve the insecticidal properties of baculovirus. Bioassays were performed to evaluate the *egt* mutants *in vivo.* Although we obtained different *egt* mutants (Table 2), the bioassay was restricted to the mutant *egt*#2 (AcMNPV-Δ49egt), which introduced an early stop codon and a frameshift mutation and to the wild-type (AcMNPV-WT) as control. At the tested concentration, no differences in mortality (pathogenicity) was observed between both viruses (Fig. 4), however, as expected from previous EGT mutants (20, 21) the virulence (time needed to kill the insects) was slightly higher for the AcMNPV-Δ49egt (Log-rank Mantel-Cox test; P<0.005). Under our experimental conditions, the Median survival time values of AcMNPV-Δ49egt (120 h) was about 10% lower than the obtained with the wild-type AcMNPV (132 h).

**Figure 4.**
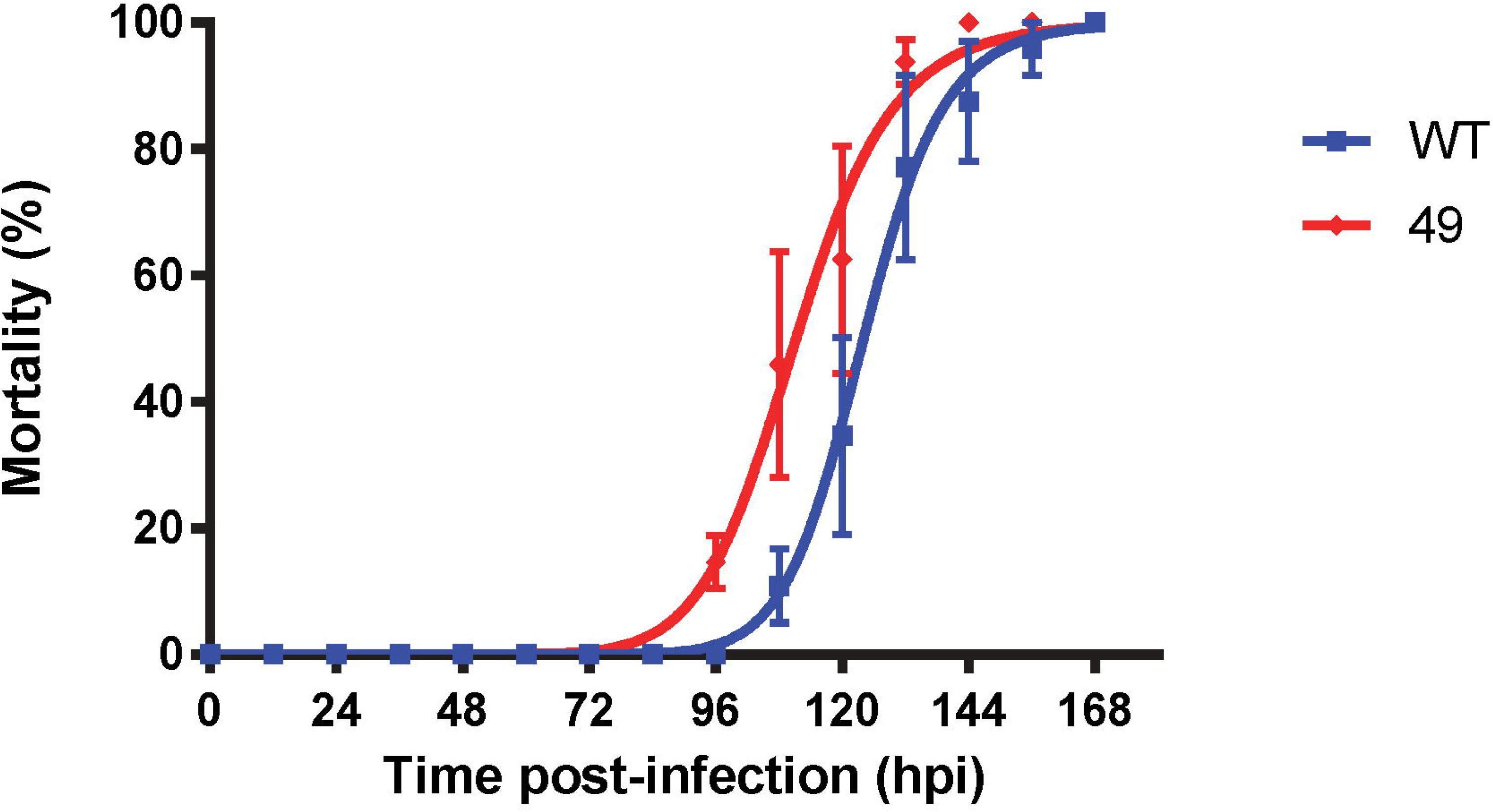
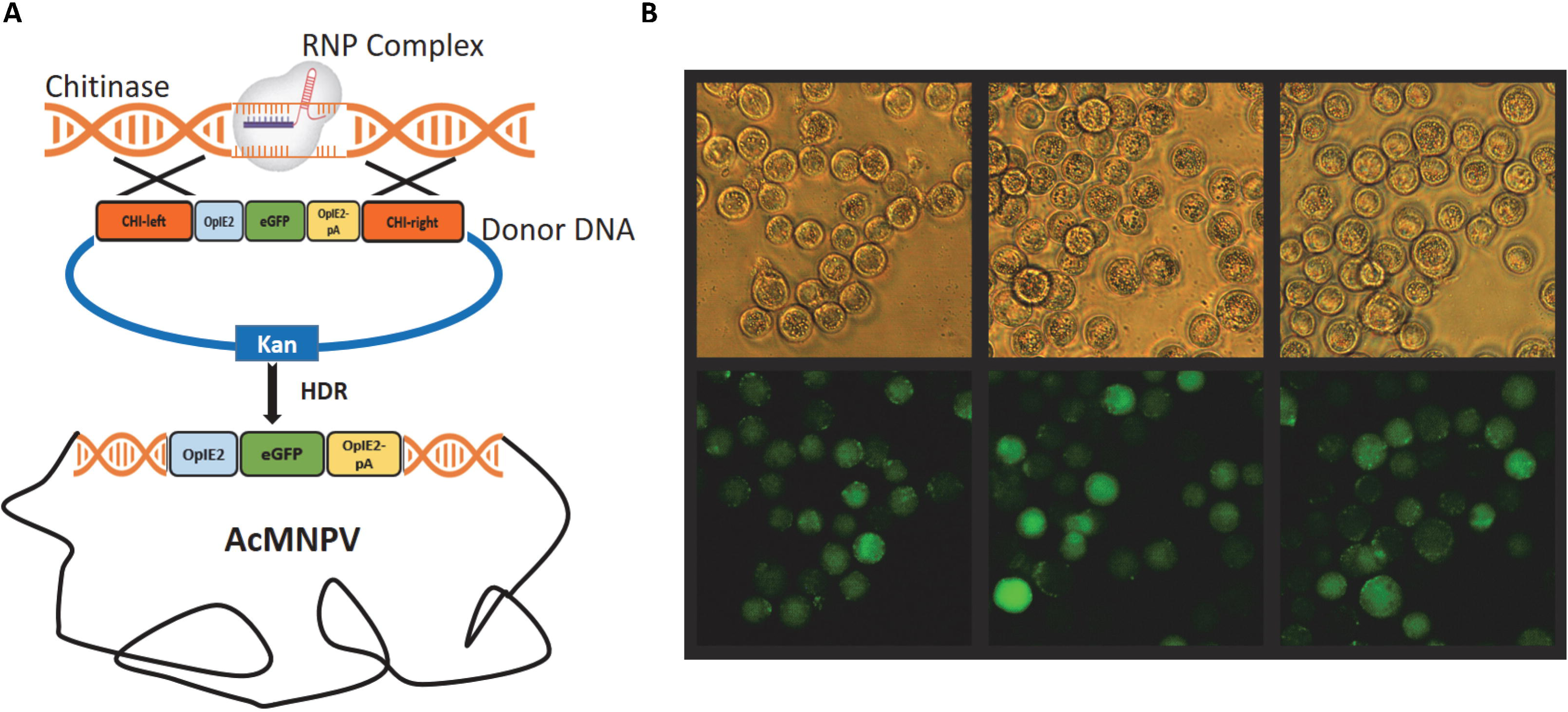
Effect of *egt* deletion on AcMNPV virulence. Time-mortality curves of the AcMNPV-WT *vs* AcMNPV-Δ49egt viruses in *S. exigua* third instar larvae. The error bars represent the standard error of the mean from 3 independent replicates.

### Gene Knock-in of wild type viruses

The capacity of our procedure to introduce foreign DNA sequences into specific regions of the AcMNPV genome was also tested. A fragment of DNA containing the *OpIE2* promoter and terminator driving the expression of the e*GFP* gene and flanked by sequences from the AcMNPV *chitinase* was used as a donor vector (Fig. 5A). The construction was then inserted into the *chitinase* locus of a recombinant AcMNPV expressing polyhedrin under the *polh* promoter (Bac-to-Bac system) using a similar procedure (see material and methods). Individual viruses obtained after the transfection were isolated by plaque assay and the rate of recombination was assessed by direct observation of eGFP production after infection of Sf21 cells (Fig. 5B). On average, 20% (4 out of 20) of the isolated clones were expressing GFP. In addition, all the GFP positive clones were simultaneously expressing the recombinant *polyhedrin* as reflected on the presence of viral OBs, indicating that CRISPR/Cas9-mediated edition does not interfere with previous transgenes from the targeted virus.

## DISCUSSION

In this study, we have shown the potential of using the CRISPR/Cas9 technology for the edition of baculovirus genomes to improve its use as protein expression system as well as insecticidal agent. Gene edition using CRISPR/Cas9 system has been recently reported in other viral genomes with biomedical implications such adenovirus (ADV) (22), type I herpes simplex virus (HSV-1) (15, 23), HIV-1 provirus (24), Epstein-Barr virus (25) and Human Cytomegalovirus (26). Our procedure has shown that baculovirus can also be edit with this system showing relatively high efficiency in the generation of random deletions or insertions in all targeted genes as well as specific gene insertions on the targeted loci.

Our procedure has shown efficient edition of all the genes that were initially targeted, though the efficiency was variable and ranged from 10 to 40% depending of the targeted gene. Since the transfection reactions were not systematically replicated, observed variability could be just caused by the stochastic variation or other factors such as the targeted sequence and the selected sgRNA (23). Although all the used sgRNA were validated *in vitro* and showed equivalent efficiency, the *in vivo* efficiency could depend on other cellular factors associated to the viral replication and the cellular repair after cleavage. It has been shown that *in vivo* efficiency can be improved using dual or multiplex sgRNA strategy (27). It would be interesting to test if the simultaneous use of multiple sgRNA targeting the gene of interest could increase the efficiency in the edition of baculovirus.

One of the advantages of the CRISPR/Cas9 technique lies in the existence of many alternative strategies for the delivering of the components into the nucleus (28). In our study, we transfected the RNP and viral DNA complex together into cells and isolated the initial viral progeny. This strategy has some advantages that could have contributed to the successful edition. First, RNP complex has easy access to the viral DNA due to simultaneous entry into the nucleus. Second, the risk of generating off-target mutations is reduced because ribonucleoprotein complexes have short half-life in the nucleus (29–31). In addition, in case of viral fitness-cost associated to the introduced mutations, the isolation of the initial progeny prevents further dilutions of the generated mutants with the non-edited genotypes.

In this study, we exploited the CRISPR/Cas9 system to engineering AcMNPV genome for different applications. We knocked-out non-essential genes for viral survival or infectivity in cell culture aiming to increase the production of recombinant proteins. We have shown that knock-out of the *ODV-E26* gene, although does not enhances protein production in cell culture, had an important increase in GFP expression in *S. exigua* larvae. The molecular function of the ODV-E26 protein is unknown, but it has been shown to interacts with the IE0 and IE1 transcription factors of AcMNPV (4, 32). These two factors are fundamental for the replication of the virus since they are the main regulators and activators of transcription. It was described that the binding sites for IE0 and IE1 in ODV-E26 are between amino acids 126-153 and 72-99 respectively. It was hypothesized that the function of ODV-E26 could be the regulation of these 2 factors, and as a consequence of ODV-E26 knock-out, the IE0 and IE1 levels increase, contributing to the increase in expression of certain genes. In the present work the RNP produces a cut-off between amino acid 23 and 24 and although some of the clones obtained had an early stop codon or a frameshift mutation that affects the presence of the, the IE0 and IE1 binding sites. However, ODV-E26#1 is not directly affected on the IE0 and IE1 binding sites and shows a significant increase in protein production in the larvae.

The other knocked-out genes did not show effect on the protein expression in cells or larvae. Nevertheless, our results have confirmed that their function is not essential for viral replication of the virus and open the possibility of using our procedure to test for simultaneous knocks-out of these or additional non-essential genes for the enhancement in the protein production. Previous studies have shown that single, as well as multiple deletion of certain viral genes, can increase the production of recombinant protein without affecting the cell viability. Different studies have shown that deletion of the *v-cath* and *chiA* genes, two enzymes responsible for the liquefaction of infected larvae improve the expression and stability of the recombinant proteins (11, 33–35). Latter improvements of these genomes were obtained by additional knocks-out of other non-essential genes for *in vitro* replication as *p26, p10* and *p74* (12). Due to the simplicity of the method reported here, procedures for systematic knock-out of all the AcMNPV predicted loci could reveal additional know-outs with enhanced protein productions.

Alternative strategy for the enhancement of protein production with baculovirus have been focused on the addition of certain genes into the viral genome. An example is the introduction of *vankyrin* genes from an insect virus *Campoletis sonorensis ichnovirus* (Fath-Goodin et al. 2009) that delays lysis of the infected cells, and an increased time for the expression of recombinant proteins. In another hand, glycosylation performance has also been improved by the simultaneous expression of glycosyltransferases (36, 37). Given the relatively high success rate obtained the gene knock-in presented in this work, our procedure could be easily applied for the simultaneous or serial insertion of those genes enhancing the expression, stability or properties of the recombinant proteins.

We have additionally shown the application of our procedure for the edition of wild type viruses and the generation of non-GM (genetically modified) viruses with potential applications in pest control. As a prove of concept we have targeted the *egt* locus. Previous studies have revealed the potential of truncation of this gene for the improvement of the insecticidal properties of baculovirus (10, 21, 38–40). Our *egt* knock-out have shown similar levels of improvement as previous studies with other *egt* knock-out mutants generated by insertion on the *egt* locus of a selecting marker (10, 21, 39) and consequently considered as GM virus. From an applied point of view, and although certain countries are currently debating about the consideration of CRISPR/Cas9 mutants as GM or not (41, 42), our results provide new tools for the generation of GM-free virus with improved properties and potential use in the field. This methodology could be extended to other viruses and other viral loci for the generation of new viral strains with improved properties. For instance, two independent isolates from the *Cydia pomonella* granulovirus (CpGV) that are able to overcome resistance in the codling moth to the classical strains of CpGV have been recently isolated (43, 44). Although the mutations on these new isolates responsible for the overcoming resistant phenotype are still unknown, gene edition using CRISPR/Cas9 approaches could be used to engineering and improvement of new viral strains. In summary, with this work we have shown how CRISPR/Cas9 methodology can also be applied to the edition of baculovirus and could contribute to the engineering of baculovirus for the increasing the protein yield and properties of the recombinant proteins expressed using the BEVS as well as a to improve the biological properties of baculovirus as a pest control agent.

## MATERIALS AND METHODS

### Cell culture and insects

The *Spodoptera frugiperda* (Sf21) insect cell line was cultured in Gibco® Grace’s Medium (1X) (Thermo Fisher Scientific, Waltham) supplemented with 10% heat-inactivated fetal bovine serum (FBS) at 25°C.

*Spodoptera exigua* larvae were obtained from our laboratory colony. Larvae were reared on an artificial diet at 25-27°C with 70% relative humidity and a photoperiod of 16/8 hours (light/dark).

### Viruses and viral DNA

Two types of AcMNPV genomes were targeted in this study. Wild type AcMNPV genome was derived from the C6 strain. The virus was replicated in Sf21 cells under standard conditions and the viral DNA was extracted using standard phenol/chloroform protocol (45). Briefly, Passage 1 (P1) viral stock was harvested and centrifuged at 500 × g for 5 min. BVs clarified were concentrated by centrifugation at 40.000 × g for 1 hour. Pellet was resuspended in 200 μl Virus Lysis Buffer (10 mM Tris-HCl pH 7.6, 10 mM EDTA, 0.5 % SDS) and 10 μl proteinase K (10 mg/ml) and incubated at 50 °C for 1 hour with shaking. 200 μl of phenol/chloroform was added and centrifugated at 14.000 × *g* for 10 min. The aqueous phase was transferred to another tube and a 1/10 of volume of 3 M sodium acetate solution, 2 volumes of absolute ethanol, and 5 μl, of glycogen (5 mg/ml) were added and kept overnight at −20 °C. The solution was then centrifuged at 14.000 × g for 15 minutes and the pellet was washed twice with ethanol 70 % by centrifuged at 8.000 × g for 5 minutes. The dry pellet was resuspended in 30 μl of TE (10 mM Tris-HCl pH 8.0, 1mM EDTA).

In another hand, the bacmid DNA from recombinant AcMNPVs expressing eGFP under *pSeL* promoter (46) or Polyhedrin under the *polh* promoter (47) were generated in previous studies of the group using the Bac-to-Bac system (Thermo Fisher Scientific). Bacmid DNA was isolated following the manufacturer’s instructions.

### AcMNPV knock-out

The following genes were targeted in the recombinant pSeL-GFP-AcMNPV baculovirus: *ODV/E26* (NC_001623.1: 13092-13769), *F-Protein* (NC_001623.1:18513-20585), *p74* (NC_001623.1:c121072-119135), and *Ac18* (NC_001623.1:c15459-14398). For the edition of the wild type AcMNPV, *egt* gene (NC_001623.1: 11426-12946) was selected.

The ribonucleoprotein (RNP) complex consisting in the Alt-R *S. pyogenes* Cas9 nuclease in complex with Alt-R CRISPR-Cas9 guide RNA (sgRNA) (IDT-Europe, Leuven) was generated *in vitro*. The sgRNA was generated by combining of the crRNA (CRISPR RNA) with the tracrRNA (trans-activating crRNA) in a 1:1 proportion, being the crRNAs the specific part of each targeted genes. The crRNAs were designed using the CHOP-CHOP online platform (https://chopchop.rc.fas.harvard.edu) and synthetized by Integrated DNA Technologies, Inc. (IDT-Europe). Gene specific crRNAs forming sgRNAs with the highest on-target and lowest off-target score were selected (Table 1). The targeted regions for each gene were limited to the 5’ region of the coding region (limited to the first 30 % nucleotides).

For the assembly of the sgRNA, 1 μl of each crRNA (1 μM) were incubated at 95 °C for 5 minutes with 1 μl of Alt-R tracrRNA (1 μM) (IDT-Europe) and 98 μl of Nuclease-Free Duplex Buffer (IDT-Europe). The RNP complex was then assembled in a final volume of 10 μl by combining 1 μl of sgRNA (100 ng/μl), 6.2 μl of the Cas9 (1 mM), and 2.8 μl of nuclease free water. Finally, the mix was incubated at 37 °C for 5 minutes and stored at 4 °C. Efficacy of each RNP complex was previously analyzed using *in vitro* Cas9 digestion (data not shown). One microgram of purified viral DNA was co-transfected with 10 μl of the RNP complex *Sf21* cells at 70% confluence in Grace’s Medium without FBS by using Cellfectin® II Reagent (Invitrogen). After 5 hours of incubation at 27 °C, the medium was replaced by Grace’s medium supplemented with 10 % FBS and incubated for 72 hours at 27 °C. The transfection efficiency was confirmed by microscopic observation of GFP (for the pSeL-GFP-AcMNPV) or the presence of viral occlusion bodies (for the wt-AcMNPV). The supernatant containing the virus was cleaned by centrifugation at 500 × g and then stored at 4 °C.

Individual viruses derived from each transfection event were isolated from the supernatants (containing a mixture of edited and non-edited viruses) by plaque assay (O’Reilly et al, 1994). The individual clones were amplified in *Sf21* cells and a fraction (50 μl) was treated with PrepMan® Ultra reagent (Applied Biosystems™, Foster city) and used for PCR amplification of the targeted region using specific primers (Table 3). PCR products were purified and sequenced by Sanger sequencing to confirm the presence of mutations in the PAM-flaking region. Edited baculoviruses were further amplified to high-titer stocks in Sf21 cells. Viral stock used in subsequent studies were tittered by qPCR using specific primers for the viral *DNApol* (Supplementary Table 1) as previously described (46). Non-edited baculovirus clones were also included as a control.

**Table 3.**
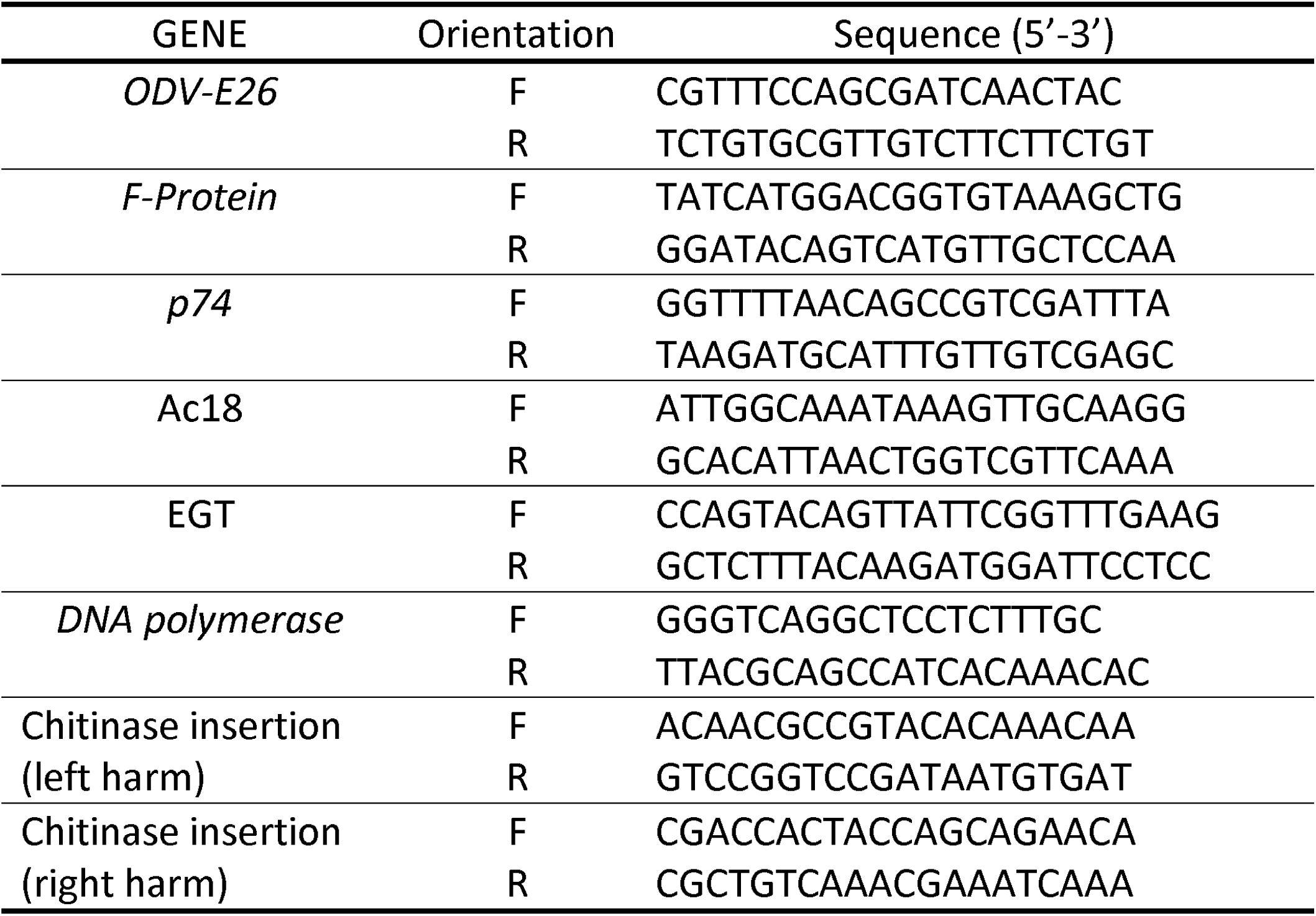
Summary of Primers employed in the study

### AcMNPV knock-in

The pU57-Kan-Chi-eGFP donor plasmid (Fig. 5) was designed using SnapGene software (from GSL Biotech; available at www.snapgene.com) and artificially synthesized by Genscript (Piscataway, NJ, USA). The length of upstream and downstream homology arms were 500 bp long and targeted the *chitinase* gene from AcMNPV. The *eGFP* gene was cloned under the control of the early viral promoter *OpIE-2*. The total length of the inserted sequence between the two homology arms was 1539 bp.

The RNP complex was generated as described above. One microgram of Viral DNA (AcMNPV expressing Polyhedrin under the *polh* promoter), 400 ng of the donor plasmid and 10 μl of the RNP complex was co-transfect in *Sf21* cells at 70 % confluence in Grace’s Medium without FBS by using Cellfectin® II Reagent. For that, viral DNA and the donor plasmid were previously mixed with the Cellfectin® II Reagent for 20 minutes, then the RNP was added and incubated for additional 20 minutes. Seventy-two hours post-transfection, the supernatant containing the virus was harvested and used to infect new Sf21 cells to confirm the presence of GFP foci (representing knock-in viruses). Individual clones were further isolated by plaque assay and amplified in *Sf21* cells. Correct location of the gene insertion on the chitinase locus was confirmed by PCR amplification and sequencing of the flanking regions (Table 3).

### Infection assay in Sf21 cell culture and *S. exigua* larvae

Sf21 cells were cultured in 24-well plate at a confluence of 70 %, then the cells were infected with the different baculoviruses at multiplicity of infection (MOI) of 0.5 and 5 and incubated at 27 °C. The cells were collected 72 hours post-infection by centrifugation at 500 × g for 5 minutes and kept at −20 °C until the quantification of GFP fluorescence expression.

*S. exigua* larvae were infected by intrahemocoelic injection with 5 μl of MQ-water containing 5 × 10^4^ BVs of the different baculoviruses (15 larvae for each assay). Larvae were incubated at 25 °C for 72 h, then were frozen at −20 °C until its analysis.

### Analysis of GFP expression

Frozen cells were lysated in lysis buffer (50 mM Tris-HCl pH 7.5, 100 mM NaCl, 1 mM DTT, 5 % glycerol) and centrifuged at 16000 × g for 1 minutes. The supernatants were used for the determination of the expression of GFP by the determination of the emission of GFP fluorescence in a microplate reader (TECAN infinite M200Pro) (λ excitation 485 nm and λ emission 535 nm). Values were expressed as the relative GFP fluorescence intensity, taken as 1 of the value corresponding to the maximum intensity obtained with the control virus after normalization to the total number of cells. For all of the experiments, the reported values correspond to at least three independent replicates.

Frozen larvae were homogenised in group of 5 larvae in extraction buffer (0,01 % IGEPAL® CA-630 (Sigma-Aldrich, Saint Louis), 1 mM PMSF, 25 mM DTT in PBS 1X). Homogenates were centrifuged at 10,000 × g for 20 minutes at 4 °C. Supernatants were diluted and used for the determination of the expression of GFP as described above. The values were normalized according to the total protein quantified by Bradford using the Quick Start™ Bradford reagent (BioRad, Hercules, CA, USA). Values were expressed as the relative GFP fluorescence intensity, taken as 1 of the value corresponding to the maximum intensity obtained with the larvae infected with the control virus.

### *In vivo* evaluation of AcMNPV-Δegt mutant

For the production of viral OBs, The *Sf21* cells culture (6×10^6^ cells) were infected with wild-type AcMNPV (strain C6) and *egt* edited virus at MOI 1 until most of cells were lysated. The resulting OBs and remaining cells were collected by centrifugation at 1,000 × g for 5 minutes and resuspension in 0,1 % SDS. The cell lysates containing the viral OBs were loaded over 40 % sucrose solution and centrifuged at 30,000 × g for 30 minutes, the pellet was washed with water and centrifuged at 2,000 × g for 5 minutes. Finally, the pellet OBs were suspended in water and quantified using a Neubauer chamber.

Bioassays were performed in third instar *S. exigua* using the droplet feeding method (48). Stock suspensions of viral polyhedral (2×10^8^ OBs/ml) were diluted, in a solution of 10 % sucrose and 1 % phenol red stain. Larvae were reared at 26 °C and mortality was recorded each 12 hours until all larvae had died. The bioassay was performed side-by-side for the two viruses (AcMNPV-Δegt and wt AcMNPV) and independently repeated three times. Mortality curves were assessed using the Kaplan–Meier method, and compared using the log□rank analysis (Mantel–Cox test) using GraphPad Prism software (GraphPad Software Inc., USA).

## Acknowledgments

We thank Rosa Maria González□Martínez for her excellent help with insect rearing and laboratory management. This work was partially supported by ERC Consolidator Grant 724519 Vis-a-Vis and the he Spanish Ministry of Economy, Industry and Competitiveness and by European FEDER funds (grant AGL2014-57752-C2-2R).

